# *In silico Drosophila Patient Model* Reveals Optimal Combinatorial Therapies for Colorectal Cancer

**DOI:** 10.1101/2020.08.31.274829

**Authors:** Mahnoor Naseer Gondal, Rida Nasir Butt, Osama Shiraz Shah, Zainab Nasir, Risham Hussain, Huma Khawar, Muhammad Tariq, Amir Faisal, Safee Ullah Chaudhary

## Abstract

*In silico* models of biomolecular regulation in cancer, annotated with patient-specific gene expression data can aid in the development of novel personalized cancer therapeutics strategies. *Drosophila melanogaster* is a well-established animal model that is increasingly being employed to evaluate preclinical personalized cancer therapies. Here, we report five Boolean network models of biomolecular regulation in cells lining the *Drosophila* midgut epithelium and annotate them with patient-specific mutation data to develop an *in silico Drosophila Patient Model* (DPM). The network models were validated against cell-type-specific RNA-seq gene expression data from the FlyGut-*seq* database and through three literature-based case studies on colorectal cancer. The results obtained from the study help elucidate cell fate evolution in colorectal tumorigenesis, validate cytotoxicity of nine FDA-approved cancer drugs, and devise optimal personalized drug treatment combinations. The proposed personalized therapeutics approach also helped identify synergistic combinations of chemotherapy (paclitaxel) with targeted therapies (pazopanib, or ruxolitinib) for treating colorectal cancer. In conclusion, this work provides a novel roadmap for decoding colorectal tumorigenesis and in the development of personalized cancer therapeutics through a DPM.

## Introduction

Cancer development is a multistep process that is driven by a heterogeneous combination of somatic mutations at the genetic and epigenetic levels [1,2]. Specific mutations in oncogenes [3] and tumor suppressor genes, [4,5] that result in their activation and inactivation, respectively, manifest themselves at tissue-level in the form of polyps, multi-layering, and metastasis [1,6,7]. These system-level properties resulting from heterogeneous biomolecular aberrations are also acclaimed as “*hallmarks of cancer”* [1,7]. The heterogeneity amongst individual cancer cells stems from factors such as genomic instability, clonal evolution, and variations in the microenvironment [8,9]. This fosters plasticity in cancer cells which leads to drug resistance – a leading impediment in the treatment of the disease [8–10]. As a result, despite major research initiatives and resultant advancements in decoding the molecular basis of cancer, a comprehensive treatment for the disease still alludes researchers. The limited therapeutic regimens approved by the Food and Drug Administration (FDA) [11–13] exhibit variable efficacies across patients besides a multitude of toxic side effects and, multi-drug resistance [14].

Towards designing efficacious personalized cancer therapeutics, recent advances in high-throughput omics-based approaches complemented by patient-specific gene expression data can provide significant assistance [15,16]. Several online databases and portals provide such freely available datasets including cBioPortal [17], The Cancer Genome Atlas (TCGA) [18], and International Cancer Genome Consortium (ICGC) [19] amongst others [20–22]. However, effective and seamless utilization of such patient-specific genomic data to design personalized cancer therapies is still a fledgling area. Researchers are increasingly employing whole-animal models [23–26] such as mouse, zebrafish, and fruit fly for preclinical *in vivo* validation of therapeutic hypotheses generated from personalized therapeutics studies. Amongst the animal models, *Drosophila melanogaster* has become a popular platform for gene manipulation, investigating site-specific changes in the genome, and high-throughput whole-animal screening [15,27]. Importantly, a comparative study of human and fly genome showed that 60% of disease causing genes in humans are conserved in *Drosophila* [28,29]. Additionally, ease of handling and significantly lower genetic redundancy imparts further advantage to the employment of fly models [5]. As a result, over 50 different data repositories, and tools are now available for hosting data on the fly genome, RNAi screens, and expression data including *Flybase* [30], FlyGut*-seq* [31], *FlyAtlas* [32] databases. Specifically in the case of cancer, several *in vivo* studies have been designed to elicit novel therapeutic targets using *Drosophila* model system [33–37]. One salient example is the validation of indomethacin, which is reported to enhance human Adenomatous Polyposis Coli (*hAPC*) induced phenotype in *Drosophila* eye [38] and therefore, employed for treating colorectal cancer (CRC). Vandetanib, another approved targeted therapy that was also validated by using *Drosophila* system, suppressed Ret activity, and was later approved for medullary thyroid carcinoma (MTC) [33,34].

However, a major shortcoming of using such mono-therapeutic agents for cancer treatment stems from the tumor heterogeneity which results in the selection of resistant cells [39,40] besides acting specifically on singular pathways. To overcome these issues, multiple therapeutic agents acting on multiple pathways in synergy can significantly increase drug efficacy, besides lowering the therapeutic dosage [40]. To evaluate potential high-efficacy synergistic drug combinations, researchers have employed *Drosophila* model in preclinical studies to elicit optimal drug combinations [36,37]. The *Drosophila* Lung Cancer Model by Levine *et al*. [36] helped identify trametinib and fluvastatin as combinatorial drug therapy for lung cancer. Further, an EGFR induced lung tumor model was also designed in *Drosophila* which assisted in providing an alternative combination of drugs for lung cancer treatment through screening an FDA-approved compound library [37]. However, combinatorial therapies pose unique challenges such as multidrug resistance in chemotherapy [14] and cross drug resistance [41,42] besides the continuing need for higher therapeutic efficacies [43]. Towards tackling these issues, researchers are now ‘personalizing’ live animal platforms for employment in preclinical studies to design efficacious therapeutic regimens. For instance, a comprehensive state-of-the-art *in vivo Drosophila Patient Model* using a personalized therapeutics approach was described in flies [44]. This particular study involved genetic manipulation of the fly genome to induce mutations specific to KRAS-mutant metastatic colorectal cancer. Combinatorial therapies were then given to the transgenic flies, harbouring mutations that were identified in the patient, to discover high-efficacy synergistic drug combinations.

Here, we propose an *in silico* counterpart of the *in vivo Drosophila Patient Model* (DPM) which will facilitate in the modeling and analysis of patient-specified CRC models besides overcoming the challenges of administering combinatorial therapies in animal models [45,46]. We have constructed five biomolecular network models of cells regulating the maintenance of adult *Drosophila* midgut epithelium lining. These include multipotent intestinal stem cells (ISCs) [47–51], enteroblasts (EBs) [52], enterocytes (ECs) [53–56], enteroendocrine cells (EEs) [57] and visceral muscle (VM) cells [58]. Next, we evaluated each network’s ability to program cell fates in normal conditions as well as under minor perturbations. The networks were then subjected to three types of inputs including physiological inputs (referred to as “*normal*”), aberrant inputs such that the fly homeostatic midgut regulation is perturbed (referred to as “*stress*”), and oncogenic inputs (referred to as “*cancer”*). The cell fate outcomes under normal and cancer conditions were validated against published literature. The individual output node propensities were also validated against RNA-seq gene expression values taken from FlyGut*-seq* [31,59] database. Finally, three literature-based case studies were constructed to further validate the proposed *in silico* DPM. The first case study replicates colorectal tumorigenesis under progressive mutations using Martorell’s CRC model [60]. In the second case study, we employed Markstein *et al*.*’s* [61] model to perform therapeutic interventions to validate the cytotoxicity of nine FDA-approved drugs. Finally, in the third case study, we reproduced Bangi*’s* KRAS-mutant CRC model [44] for evaluating optimal personalized drug treatment combinations by incorporating key patient-specific mutations into our model followed by combinatorial therapeutic screening. Building on these case studies, we devised a novel synergistic combination of paclitaxel (a chemotherapeutic agent) and pazopanib, and ruxolitinib (targeted therapies) for treating ten CRC patients taken from cBioPortal [17,62]. The results obtained from combinatorial chemo- and targeted therapies show up to 100% increase in anti-cancerous cell fates such as apoptosis and a 100% reduction in tumorigenesis promoting cell fates such as hyper-proliferation.

Taken together, we have proposed a computational framework in the form of an *in silico* DPM to provide personalized CRC therapeutics. This approach can help reduce cancer treatment costs and facilitate in the development of higher efficacy combinatorial therapies for cancer as well as to elucidate novel therapeutic targets.

## Results and discussion

### Network construction and robustness analysis of regulatory homeostasis in *Drosophila melanogaster* midgut

To investigate the biomolecular signaling regulating homeostasis in *Drosophila melanogaster* midgut, we undertook an extensive literature survey and constructed five cell-type-specific rules-based network models. These models correspond to the five cellular phenotypes lining the *Drosophila* midgut which include: intestinal stem cells (ISCs) [47–51], enteroblasts (EBs) [52], enterocytes (ECs) [53–56], enteroendocrine cells (EEs) [57], and visceral muscle (VM) [58] (Tables S1-5). The scheme of pathways integration for ISC, EB, EC, EE, and VM is provided in Figures S1-5 and the resultant network models consisted of 33 nodes and 51 edges, 30 nodes and 46 edges, 24 nodes and 36 edges, 24 nodes and 36 edges, and 27 nodes and 38 edges, respectively (**Figures 1A and B**, Figures S6-10).

**Fig. 1.**
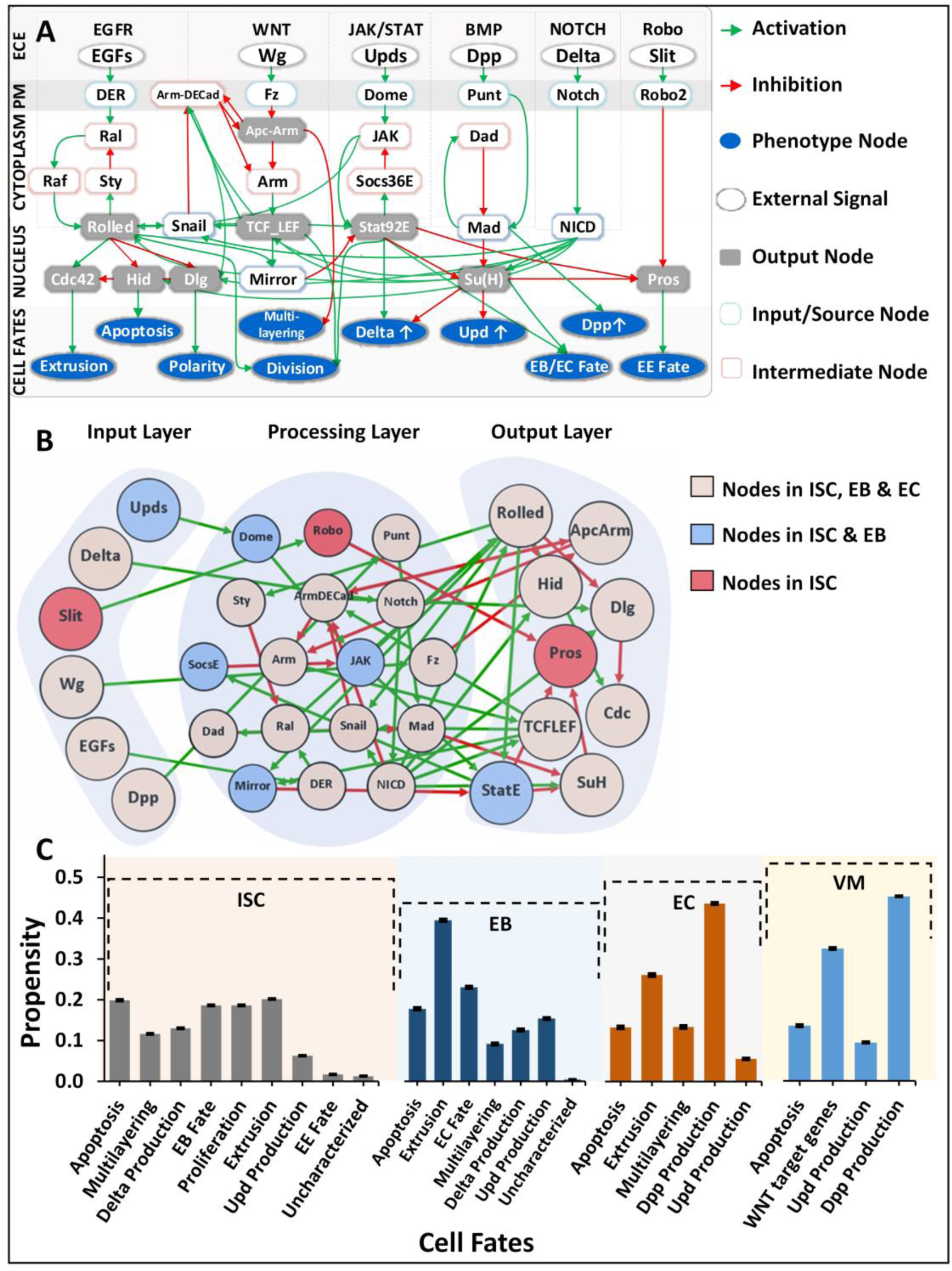
Regulatory schema of networks for the three cell types present in *Drosophila melanogaster* midgut. (**A**). The overall scheme of six conserved pathways involved in the regulation and homeostasis of an adult *Drosophila* midgut. (**B**). The mapping between input, processing, and output nodes present in the biomolecular network models of three cell types i.e. ISC, EB, and EC. (**C**). Cellular fate propensities for ISC*–*Apical, EBs, ECs, and VM, along with their respective SEMs.

Next, to evaluate the biological plausibility of the networks, we analyzed each network under normal conditions. Specifically, the biomolecular network of ISC–Apical cells exhibited extrusion, apoptosis, proliferation, and differentiation (or EB fate) with 0.182, 0.179, 0.168, and 0.168 propensities, respectively. EC network exhibited dpp production, and extrusion with corresponding propensities of 0.428, and 0.256. Lastly, for EB and VM cells, extrusion and dpp production were programmed with propensities of 0.335 and 0.450, respectively. Robustness analysis performed by inducing a 10% perturbation in the input stimuli showed the highest variations in ISC’s propensity for apoptosis (SEM=0.0014). Similarly, for EB, EC, and VM, the highest variations in propensity were observed for apoptosis with SEM=0.0027, 0.0034, and 0.0024, respectively (**Figure 1C** and Figure S11). These results indicate that all five networks are biologically plausible as they exhibited robustness against random perturbations and are hence feasible for employment in onwards analyses [63,64] (Table S6 and Figure S12).

### Evaluation and validation of biomolecular network models under normal, stress and colon cancer conditions

To evaluate the proposed networks against published literature and RNA-seq data from FlyGut*-seq* [31], Deterministic Analysis (DA) was performed [65] under normal, stress, and cancerous conditions (construed as a combination of inputs) (Table S7). Results from our analyses (**Figure 2**) revealed that in normal conditions, ISC–Apical network programmed extrusion, apoptosis, proliferation, and differentiation (or EB fate) with propensities of 0.183, 0.178, 0.168, and 0.168, respectively (Table S8). Under stress conditions, the propensity for differentiation increased to 0.213, while proliferation, and extrusion reached to 0.211, and 0.213, respectively (see Materials and methods). Lastly, in cancerous conditions, propensities for multi-layering, and apoptosis increased to 0.317, and 0.225, respectively. The results were again validated from the literature which supports elevated apoptosis in ISC’s when under extreme toxic conditions [66,67]. Literature reports also that ISCs upon encountering extreme stress, exhibit epithelium multi-layering, augmented by overgrowth [68]. Alongside, we also observed a reduction in proliferation, which corroborated with studies showing that tumor cells typically experience limited nutrient availability [69] which also slows down normal ISC cell division rate [70,71] (Figures S13-15).

**Fig. 2.**
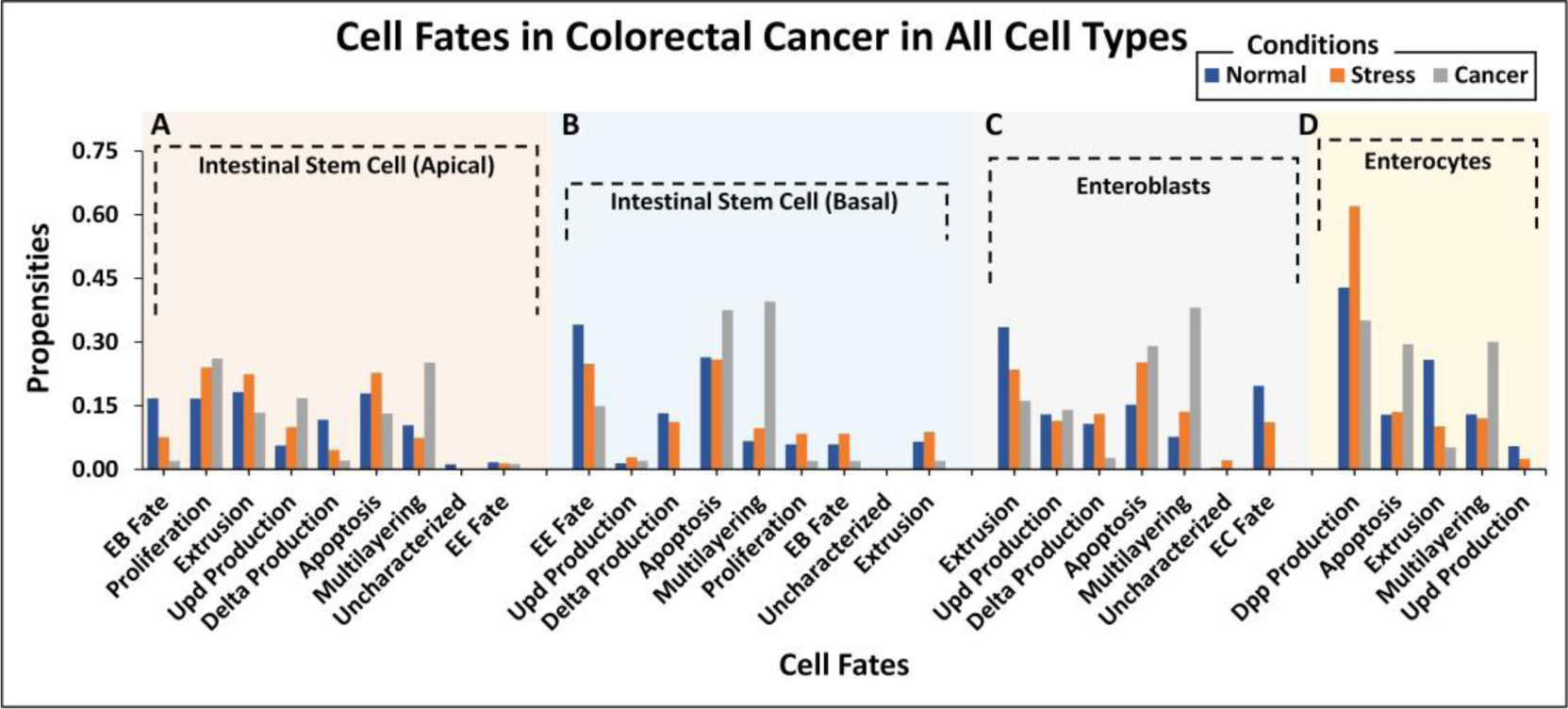
Stack bar chart representation of cell fate propensities for intestinal stem cells (ISCs) in apical and basal compartments, enteroblasts (EBs) and enterocytes (ECs) in normal, stress and cancer conditions. (A). ISC*–*Apical cells adopt nine different cell fates while one remains uncharacterized in three ambient conditions. In normal conditions, the highest propensity was observed for extrusion followed by apoptosis, proliferation, and EB fate, in order. In the case of stress, the highest propensity is that of extrusion, followed by EB fate and proliferation. In cancer, the highest propensity is that of multi-layering, followed by apoptosis and extrusion. (B). ISC*–*Basal adopts nine different cell fates with the highest propensity being for EE fate in normal conditions, apoptosis in stress conditions while in the case of cancer, multi-layering and apoptosis showed the highest propensity. (C). Seven cellular fates in EB, with the highest propensity for extrusion in normal, apoptosis in stress, and multi-layering in cancer. (D). Five cellular fates in EC, with the highest propensity for dpp production in normal, stress and cancer conditions.

For network regulation of ISC–Basal cells in physiological conditions (Table S7), the cell fate outcomes included differentiation (or EE fate), apoptosis and delta production, with propensities of 0.341, 0.264, and 0.132, respectively (Table S8). Under stress, Upd production fate increased (from 0.014 to 0.028), and differentiation rate decreased to 0.248. However, delta production remained steady. For cancer conditions, the propensity of differentiation and proliferation decreased to 0.149 and 0.020, respectively, whereas both apoptosis and multi-layering increased to 0.375 and 0.395, respectively. Both of these results, along with the relatively negligible delta expression, are in accordance with previously published reports. Moreover, extreme cellular environments are known to increase apoptosis rate in Enterocytes [66], suggesting that in absence of mutations, normal cells residing in toxic and oncogenic environments can be stressed leading to high apoptosis rates along with an inhibition of cell proliferation (Figures S16-18).

Next, we evaluated cell fate programming of the EB network under normal conditions (Table S7). The results showed extrusion, differentiation (or EC fate) and apoptosis cell fates with propensities of 0.335, 0.197, and 0.152, respectively (Table S8). Alongside, Upd production was also observed with a propensity of 0.130. However, in stress conditions, the propensity for apoptosis and multi-layering increased to 0.253 and 0.136, respectively, whereas, extrusion and differentiation (or EC fate) decreased to 0.235, and 0.111, respectively. In cancerous conditions, the salient cell fates programmed included multi-layering, apoptosis, and extrusion with propensities of 0.381, 0.291, and 0.161, respectively. Also, differentiation was suppressed to 0.140 due to toxic cellular environments. The trend in cell fate propensities in cancerous conditions also exhibited multi-layering [68] along with low delta production and extrusion (Figures S19-21), which corroborates with published literature which states that delta is a known marker for ISC and in case ISC proliferation, is reduced along with delta production [66]. Extrusion is triggered by over population of cells [72], however, in high stress conditions, cells preferentially inhibit proliferation followed by an enhanced apoptosis thereby limiting extrusion.

Moreover, the EC network was also analyzed for response under normal conditions (Table S7). The emergent cell fates included dpp production, extrusion, and multi-layering with propensities of 0.429, 0.258, and 0.130, respectively (Table S8). Under stress, extrusion rate decreased to 0.101, while apoptosis and dpp production increased to 0.135 and 0.620, respectively. In cancer conditions, however, an increase in propensities of multi-layering (0.378) and apoptosis (0.295) was observed which is in agreement with published studies [66,68] (Figures S22-24).

Lastly, a comparison of output node values for ISC–Apical, EB, and EC networks in normal conditions was performed against experimental RNA-seq data from the FlyGut*-seq* database [31]. Note that due to the paucity of regulatory dynamics in the literature on EE and VM cells, we could not evaluate their networks further. The output node propensities for ISC, EB, and EC were found to be comparable with values from the FlyGut*-seq* database [31] (**Figure 3** and Table S9).

**Fig. 3.**
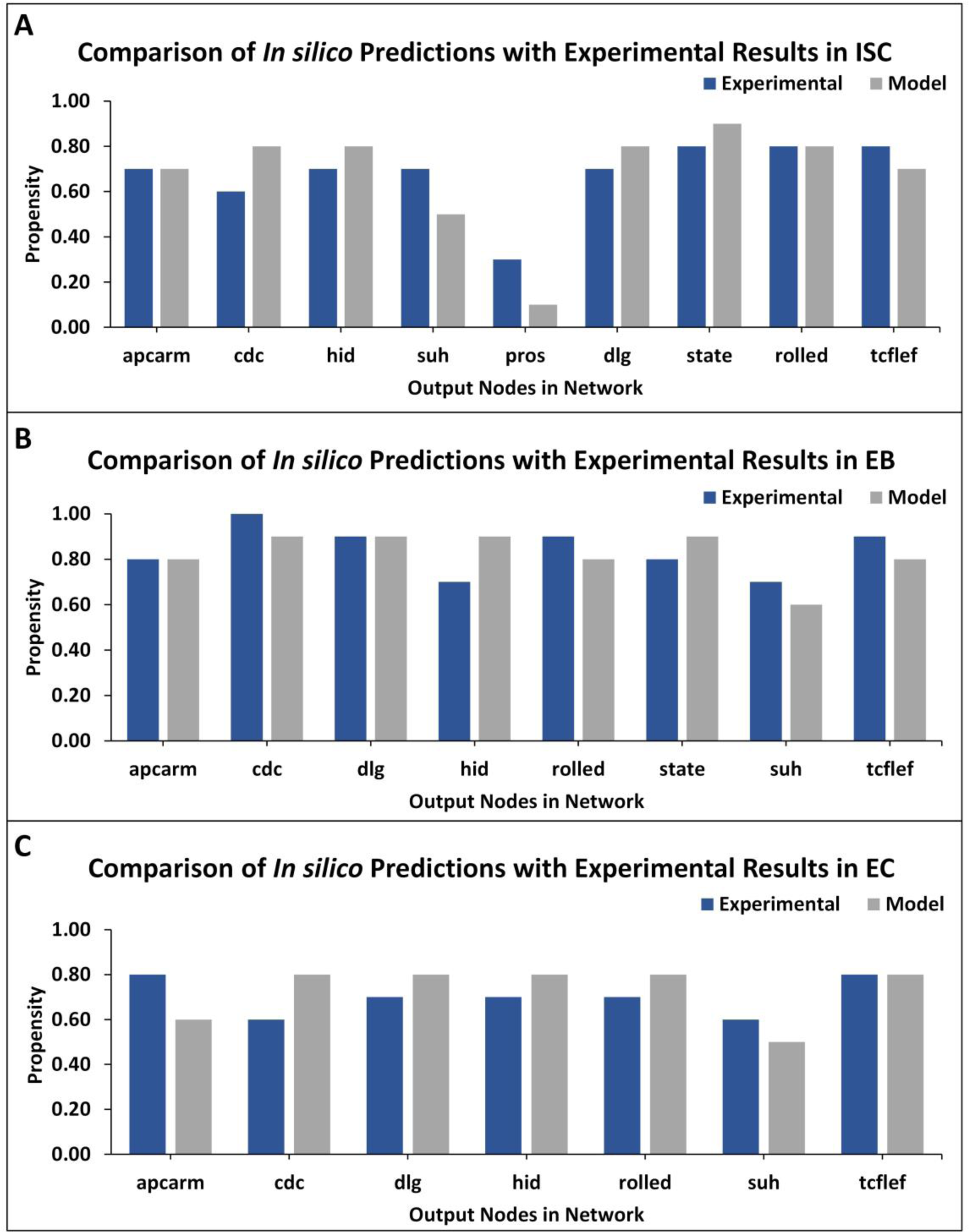
TISON output nodes propensities (*in silico* results) validation from FlyGut-*seq* database (*in vivo* results). (**A**). Comparison of nine output nodes propensities in ISC*–*Apical network: adenomatous polyposis coli (*Apc2*), cdc42 (*Cdc42*), head involution defective (*hid*), suppressor of hairless (*Su(H)*), prospero (*pros*), discs large 1 (*dlg1*), signal-transducer and activator of transcription protein at 92E (*Stat92E*), rolled (*rl*) and pangolin (*pan*). (**B**). Comparison of eight output nodes propensities in EB network: adenomatous polyposis coli (*Apc2*), cdc42 (*Cdc42*), discs large 1 (*dlg1*), head involution defective (*hid*), rolled (*rl*), signal-transducer and activator of transcription protein at 92E (*Stat92E*), suppressor of hairless (*Su(H)*), and pangolin (*pan*). (**C**). Comparison of seven output nodes propensities in EC network: adenomatous polyposis coli (*Apc2*), cdc42 (*Cdc42*), discs large 1 (*dlg1*), head involution defective (*hid*), rolled (*rl*), suppressor of hairless (*Su(H)*), and pangolin (*pan*) (Table S10).

### Case Study 1 – Investigating colorectal tumorigenesis under progressive mutations in *Drosophila* midgut

To decode the emergent cell fates during initiation and progression of colorectal cancer (CRC) in the adult *Drosophila* midgut, two salient driver mutations [60] in adenomatous polyposis coli (Apc, in WNT pathway) [73] and Ras (in the EGFR pathway) [74] were incorporated into the ISC–Apical network. These mutations were initially incorporated to act individually and later simultaneously (Figure S25). The emergent cell fates in the control case included apoptosis, proliferation, and differentiation, along with loss of polarity, multi-layering, and extrusion with propensities of 0.180, 0.168, 0.168, 0.00, 0.105 and 0.182, respectively. Upon incorporation of Apc mutation into the ISC– Apical network, a slight decrease in apoptosis and proliferation was observed as their propensities decreased to 0.165 and 0.138, respectively. Differentiation and extrusion also got reduced to 0.138 and 0.148, respectively, while multi-layering increased to 0.349, and loss of polarity remained unaffected. Next, upon introducing Ras mutation, a decrease in apoptosis (0.089) and an increase in proliferation (0.186) was observed, which indicated cellular overgrowth. Furthermore, in line with Martorell *et al*. [60], differentiation remained unchanged while the loss of polarity and extrusion increased to 0.089 and 0.255, respectively.

On the other hand, the concurrent incorporation of Apc and Ras mutations resulted in hyper-proliferation and overgrowth as apoptosis decreased to 0.066 and proliferation increased to 0.203. Differentiation rate was observed to be 0.138 and loss of polarity, multi-layering and extrusion increased to 0.066, 0.203, and 0.203, respectively. Hence, with concurrent mutations in Apc and Ras, the emergent cell fates started exhibiting the hallmarks of cancer including abnormal proliferation and loss of differentiation, etc [75]. These results were also coherent with both the experimental findings reported by Martorell *et al*. [60] (**Figure 4** and Table S11) and differential gene expression data [76] (Table S12).

**Fig. 4.**
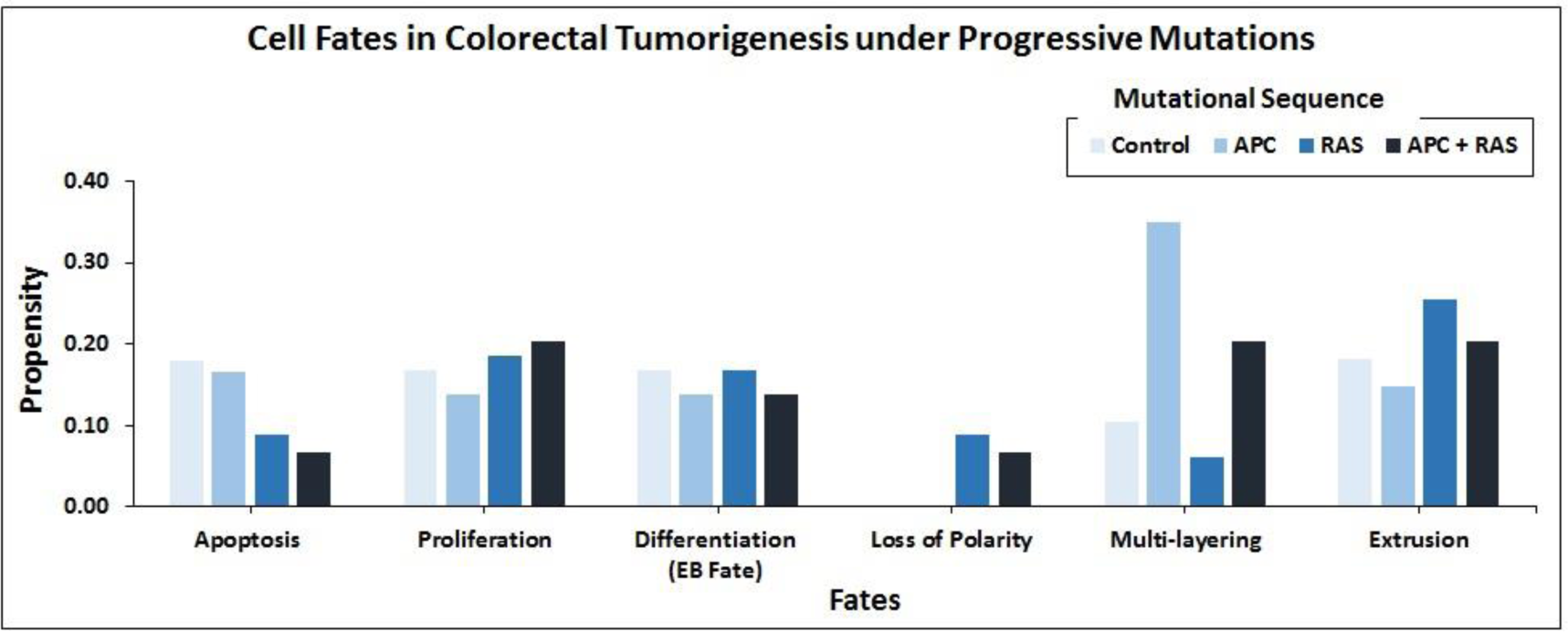
Cell fate outcomes after the introduction of progressive CRC mutations and their validation against Martorell *et al.’*s *Drosophila* CRC model. A high rate of extrusion and loss of polarity was observed in Apc-Ras as well as Ras clones. Alongside, an increased proliferation rate with a decreased apoptosis and differentiation is also highlighted by Martorell *et al*. in their *in vivo* model.

### Case Study 2 – Therapeutic evaluation of CRC in *Drosophila* midgut using targets from the literature

Introduction of gain-of-function Raf-specific driver mutations in our ISC–Apical network enabled the replication of Markstein *et al*.*’s* [61] therapeutic screen towards a comparative cytotoxicity evaluation of nine FDA-approved drugs. In their gain-of-function Raf tumor model, Markstein and colleagues had classified FDA approved drugs into class I and II drugs. According to the study class I drugs induced CRC reversal in mutated cells without effecting the wild type cells, whereas class II drugs besides reversing CRC in mutated cells, also induced CRC in wild type cells (Table S13). The result of our network analysis of the control case exhibited proliferation and apoptosis with propensities of 0.167 and 0.179, respectively. However, after the induction of Raf mutations, proliferation (0.187) rate increased along with a decrease in apoptosis (0.088). Treatment of a Raf-mutated network using class I drugs led to a decrease in proliferation (0.108) and an increase in apoptosis (0.167). No effect was observed on proliferation, which remained steady at 0.168 whereas a slight decrease was observed in apoptosis (0.167) for the wild type in comparison with the control. This confirmed the action of class I drugs which act to significantly reduce cancerous fates in CRC without having a major impact on wild type cells.

Alternatively, in the case of class II drugs, the wild type also exhibited hyper-proliferation after therapy with its propensity reaching up to 0.240 and apoptosis decreasing to 0.068. Importantly, for the CRC network, drug action continued to show cancer reversal with the propensity of proliferation around 0.108 and apoptosis at 0.132. These results suggest that class II drugs are indeed associated with drug cytotoxicity as they revert cancer phenotype but at the same time induce malignancy in normal cells under therapy. This again confirms Markstein *et al*.*’s* study which hypothesized that the extracellular environment in animal models is crucial in drug delivery and cytotoxicity (**Figure 5** and Table S14).

**Fig. 5.**
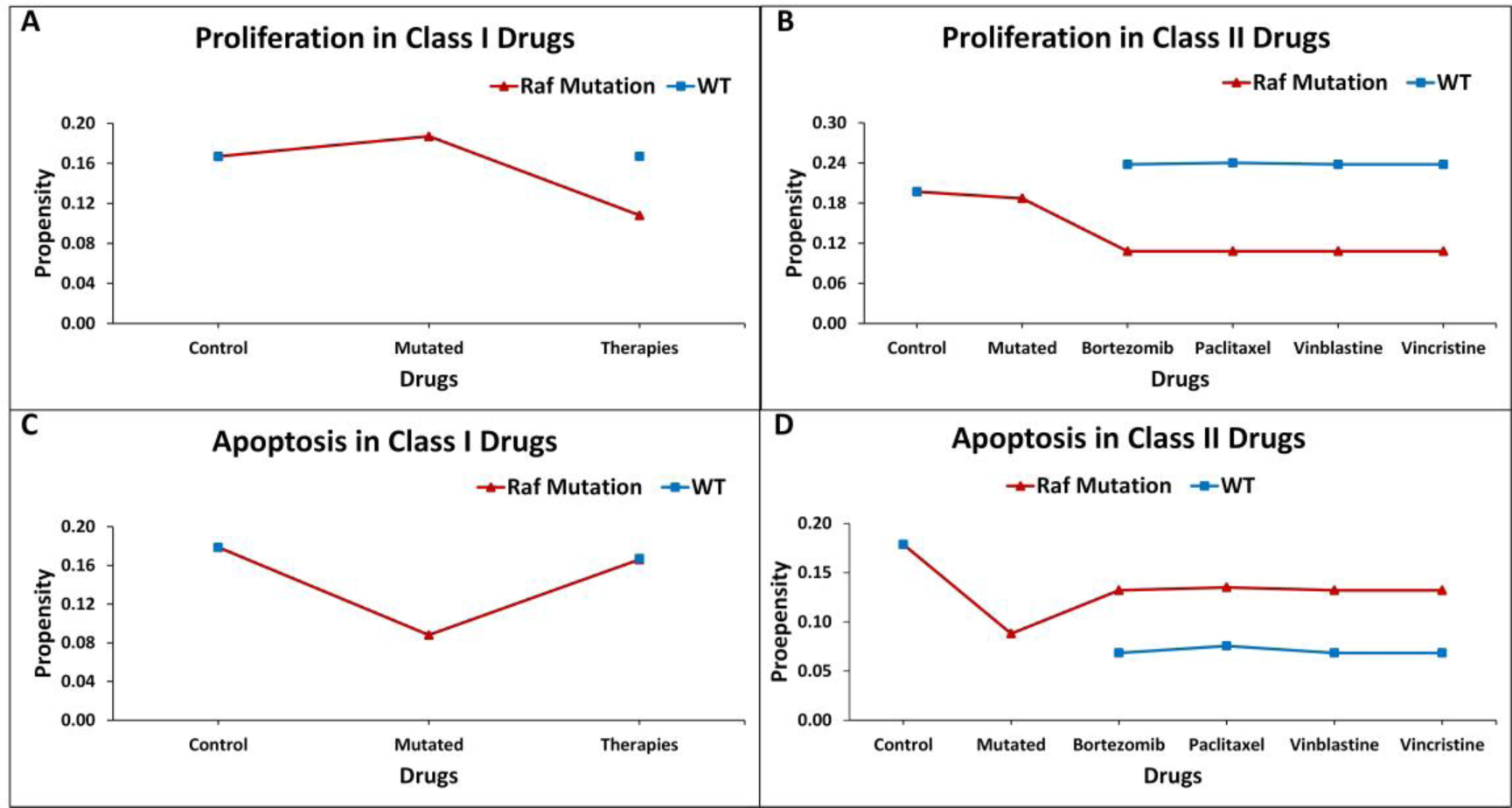
Evaluating cell fates under therapeutic screens taken from Markstein *et al*.’s *Drosophila* model. (**A**). The effect of class I drugs on cell proliferation in wild type and CRC networks, (**B**). The effect of class II drugs on apoptosis in wild type and CRC networks, (**C**). The effect of class I drugs on apoptosis in wild type and CRC mutated networks, (**D**). The effect of class II drugs on apoptosis in wild type and mutated network.

### Case Study 3 – Employing the *in silico Drosophila Patient Model* for personalized therapeutics

Towards developing a *Drosophila-based* platform for employment in orchestrating patient-centric cancer therapeutics, we adopted Bangi *et al*.’s [44] *in vivo Drosophila Patient Model* (DPM). The *in vivo* model was first translated into an *in silico* DPM which incorporated patient-specific mutations from Bangi *et al*.’s study. These mutations included eight tumor suppressors: Apc, Tp53, Fbxw7, Tgfbr2, Smarca4, Fat4, Mapk14, and Cdh1, along with one oncogenic mutation in Kras (Table S15). After inducing these patient-specific mutations into the ISC–Apical network, we administered the combinatorial therapy of trametinib and zoledronate. Our results showed that in control (i.e. healthy cells), the cell fate propensities for proliferation and apoptosis came out to be 0.167 and 0.182, respectively. Upon induction of mutations, proliferation increased to 0.250 and apoptosis decreased to 0.000, respectively. Next, with the administration of trametinib, an inhibitor of MEK kinase (mitogen-activated protein kinase kinase), used to treat patients with Kras mutation [44], the propensities for proliferation and apoptosis reverted to 0, however, upon augmentation of therapy with the addition of zoledronate in combination with trametinib, a decrease in proliferation to 0.168 and an increase in apoptosis to 0.131 was observed. These results exhibited cancer reversal on the administration of the drug combination and corroborate with Bangi *et al*.’s findings (**Figure 6)**.

**Fig. 6.**
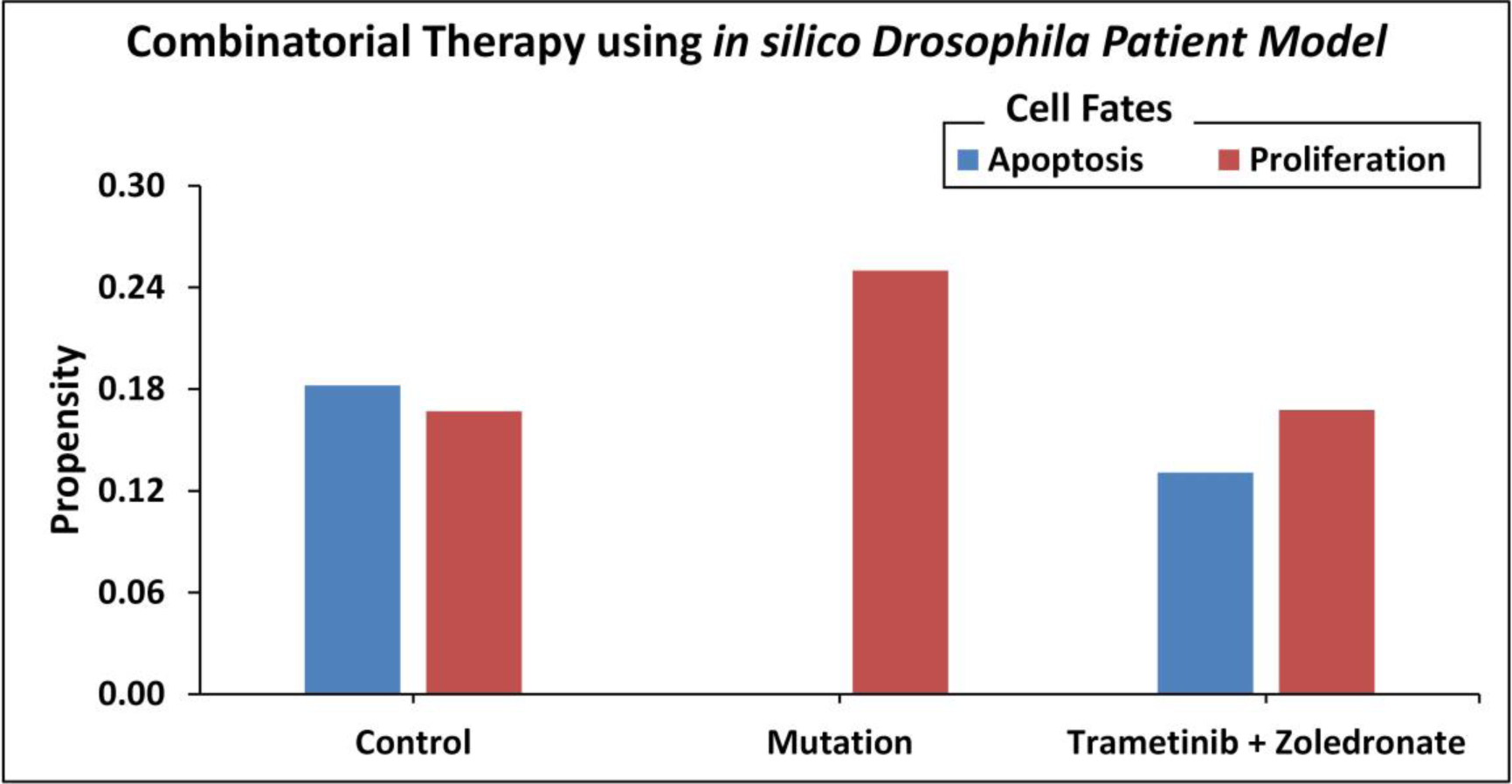
Cell fate propensities obtained from the *in vivo Drosophila* Patient Model using Bangi *et al*.’s study.

### Identification and evaluation of personalized therapeutics for CRC patients using *in silico* DPM

Towards developing personalized combinatorial therapies for treating colorectal cancer patients, we coupled our *in silico* DPM with patient-specific gene expression data from cBioPortal [17]. Patient-specific potential druggable targets were identified and their oncogenic cell fate propensities were obtained using DA pipeline. Each node was then queried in the PanDrugs database [77] to find out the drugs that targeted them directly or indirectly (Table S16). The results from this exercise elicited paclitaxel [78] and several other targeted therapies including pazopanib, and ruxolitinib depending on patient-specific mutations (Table S17). Follow up literature review showed that these drugs and their combinations are currently being used in several studies and clinical trials [79–82]. Specifically, the combination of paclitaxel and ruxolitinib was evaluated in 2018 to treat human ovarian cancer [79], while the paclitaxel-pazopanib combination was evaluated for treating metastatic melanoma [80] and is in clinical trials for Non-Small Cell Lung Cancer (NSCLC) [81] as well as angiosarcoma [82].

To test the efficacy of these drug combinations in CRC patients, we administered these therapies using the proposed *in silico* DPMs to ten patients with colorectal adenocarcinoma obtained from cBioPortal [17]. To implement the simultaneous action of chemotherapy wherein the drug introduces widespread inhibition of mitosis by stabilizing polymerized microtubules and not allowing them to function during cell division for that, we surveyed the existing literature and constructed a microtubule network (Table S18) with 23 nodes and 28 edges (Figure S26). This network was then integrated into our existing ISC-Apical network to study the behaviour of microtubule stabilization-induced cell fates in chemotherapy (Table S19). The resultant integrated network consistent of 39 nodes and 64 edges (Figure S27). Our results from combinatorial chemo- and targeted therapy using the integrated network showed up to a 100% increase in apoptosis cell fate and a 100% decrease in proliferation rate (Table S20). With administration of targeted therapy only, our results showed up to a 600% increase in apoptosis cell fate and a 100% decrease in proliferation rate (**Figure 7** and Table S21).

**Fig. 7.**
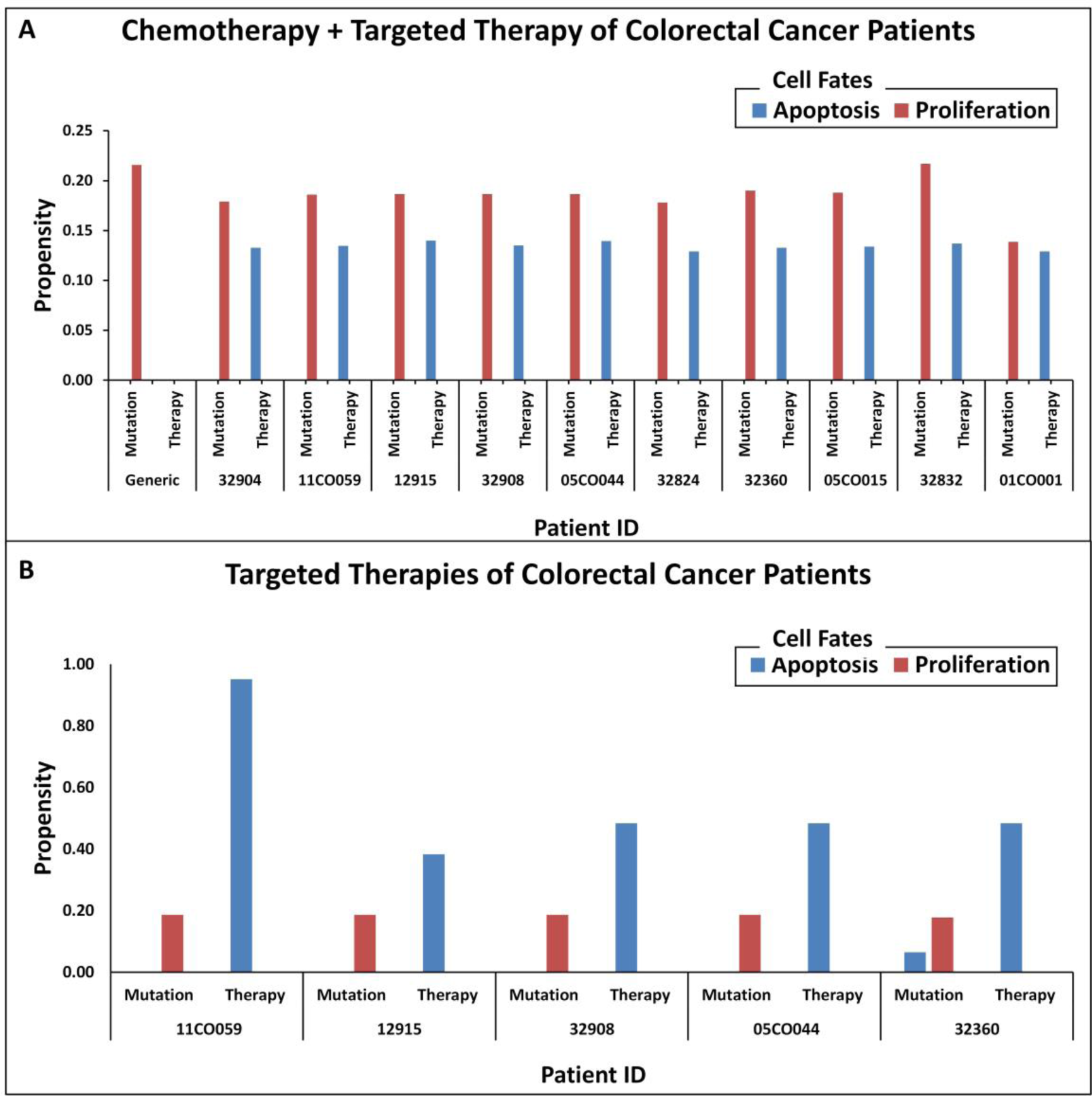
Comparison of oncogenic cell fate propensities obtained from chemotherapy + targeted therapy results versus targeted therapy results. (A) Chemotherapy + targeted therapy for ten colorectal cancer patient for personalized screening. Patient ID and mutation data were extracted from cBioPortal and cell fates for apoptosis and proliferation were plotted to observe before and after therapy results, (B). Targeted therapy for five colorectal cancer patients for personalized screening. Patient ID and mutation data were extracted from cBioPortal and cell fates for apoptosis and proliferation were plotted to observe before and after therapy results.

## Conclusion

Taken together, in our study we present a computational framework using a literature-derived *in silico Drosophila Patient Model* (DPM) for treating colorectal cancer (CRC). We carried out an extensive literature survey to construct five biomolecular network models (intestinal stem cells (ISCs) [47–51], enteroblasts (EBs) [52], enterocytes (ECs) [53–56], enteroendocrine cells (EEs) [57], and visceral muscle (VM) [58] regulating the maintenance of the epithelium in *Drosophila* midgut. The networks were analyzed in normal conditions for their robustness against minor perturbations followed by an evaluation in normal, stress, and cancer conditions. The network model was further validated against RNA-seq datasets from FlyGut-*seq* database as well as three literature-based case studies. Therapeutic screening using the proposed *in silico* DPM helped personalize treatment for individual patients taken from cBioPortal (Table S17). Outcomes from the *in silico* screening of ten patients highlight the need for a detailed evaluation of paclitaxel, a chemotherapeutic agent,and targeted therapy synergy to treat CRC patients. To the best of our knowledge, the proposed model is the first of its kind to model fly gut homeostasis and tumorigenesis using the five cells lining the midgut epithelium.

The proposed model can be deployed by wet-lab biologists in preclinical settings to evaluate potential drug targets before their *in vivo* evaluation. The flexibility offered by this model can also facilitate the incorporation of patient-specific gene expression data towards directly evaluating potential drugs. It will be interesting to employ the proposed model by investigating fly embryo formation and development by incorporating developmental genes. The *in silico* DPM further stands to strengthen the fly community by providing a tool for value addition in the development of novel therapeutic strategies using personalized therapeutics approaches.

## Materials and methods

### Data collection and Boolean modeling of five cell-type-specific networks in *Drosophila* midgut

To construct the biomolecular network models involved in cellular regulation of *Drosophila* midgut, a comprehensive review of the existing literature and databases was undertaken. The databases employed included the Kyoto Encyclopedia of Genes and Genomes (KEGG) [83], Drosophila Interactions Database (DroID) [84], and data repositories such as FlyGut*-seq* [31], and Flybase [85]. Alongside, network models of *Drosophila* by Giot *et al*. [86], Formstecher *et al*. [87], and Toku *et al*. were used to construct five rule-based Boolean biomolecular networks of the conserved signaling pathways in intestinal stem cells (ISCs) [47–51], enteroblasts (EBs) [52], enterocytes (ECs) [53–56], enteroendocrine cells (EEs) [57], and visceral muscle (VM) [58]. Six major pathways involved in maintaining the overall homeostatic nature of the fly midgut were selected from the available literature. These included Notch [88], BMP [88], EGFR [89], WNT [90], JAK-STAT [90,91], and Robo [92] pathways for each cell type lining the midgut. The network steady states were used to program cell fate outcomes such as cellular differentiation, proliferation, apoptosis, EC fate determination, etc. Boolean equations [93] were used to model the regulation of each node in the biomolecular network. TISON, an in-house theatre for *in silico* systems oncology (https://tison.lums.edu.pk) was used to translate Boolean rules into network models (see Supplementary Data).

### Robustness analysis

To validate the biological plausibility of the proposed networks, robustness analysis was performed. Physiological conditions were maintained during this process and the input node values were taken from the FlyGut*-seq* database [31,59]. The normal node states for ISC, EB, EC, and VM were perturbed by ±10%. Bootstrapping was employed on 10,000 network states. The means and standard deviations of the emergent cell fates were then calculated and standard error of means (SEM) was plotted for each cell fate to determine the biological plausibility of the scale-free networks [94] (see Supplementary Data 1).

### Deterministic analysis

The Boolean network models were analyzed using Deterministic Analysis (DA) [65,95] performed in TISON, an in-house web-based multi-scale modeling platform for *in silico* systems oncology. The DA pipeline was derived from ATLANTIS [96]. DA was used to identify ‘*cell fate attractors’* – the most probable biological states of a cell and compute their propensities. TISON’s DA pipeline requires three different input files including (i) network file, (ii) fixed node states file, and (iii) cell fate classification file. The network file contained the Boolean rules for rules-based biomolecular networks. The fixed node states file contained fixed values for generating environmental conditions such as normal, stress, or cancer conditions. The cell fate classification file was used to the map network states onto the biological cell fates in the light of particular cell fate markers [96] (Table S22). For DA, bootstrapping was employed on 10,000 network states. TISON’s *Therapeutics Editor* (TE) was used to undertake therapeutic evaluation on the network using the DA pipeline, with mutation and drug data integrated. Fixed node states for normal conditions were obtained from FlyGut-*seq* database while for cancer conditions, literature was surveyed to find out if the pathway is up or downregulated. For stress, abnormal values were abstracted by perturbing the stimuli in normal conditions (see Supplementary Data 2).

### Output node validation against Flygut-*seq* database

To validate the output node propensities of ISC-Apical, EB, and EC networks with FlyGut*-seq* database, we exported the RNA-seq, rpkm values, from the database. The dataset was then used to extract the relevant genes present in our networks (ISC, EB, and EC) using their biological names. Expression data across the five regions of the midgut (i.e. R1, R2, R3, and R5) [97] was normalized for each gene in specific cells. The normalized values were taken as normal input fixed node states for onwards analyses. The normalized values were then compared with the output node propensities from DA that was performed in normal cell fate conditions in TISON (see Supplementary Data 2).

### Cell fate data collection for case studies and their validation

To validate and exemplify our network models, we used three literature-based case studies on colorectal tumorigenesis in *Drosophila* melanogaster. For case study 1, data including cell fates under Apc and Ras single and simultaneous mutations were obtained from Martorell *et al*.’s model [60]. The differential gene expression screens and data were also obtained from Martorell *et al*. [76] (see Supplementary Data 3). TISON’s TE was used to implement the mutations in our network using TE’s horizontal therapy pipeline. For case study 2, therapeutic screens including them existing list of FDA-approved drugs for targeting ISC in *Drosophila* were adapted from Markstein *et al*.*’s* [61] study. Existing databases on drugs and drug-gene interactions such as PharmacoDB [98], PanDrugs [77], and DGIdb [99], etc [100,101] were then used to identify nodes in our ISC–Apical network, which were targets of the drugs mentioned in Markstein *et al*.*’s* study. TE was employed to deliver drug data into the CRC mutated network using TE’s vertical therapy pipeline. Different fixed node states and cell fate classification files were employed to perform network evaluations in wild type and CRC to mimic different cellular environments in normal and cancerous conditions (see Supplementary Data 4). For case study 3, patient-specific mutations, along with combination therapy drug candidates were taken from Bangi *et al*.’s [44] study. Drug databases were used to identify nodes in the ISC–Apical network which were targets of drugs mentioned in Bangi *et al*.’s study. Drugs which did not have direct targets in the network were implemented indirectly using literature-based mechanisms (see Supplementary Data 5).

### Development of an *in silico Drosophila Patient Model* (DPM) and its validation

Towards devising a novel drug combination for the treatment of colorectal tumorigenesis, we performed an exhaustive evaluation of each node in our ISC– Apical network using TISON’s TE therapy panel. For that, we started with the sensitivity analysis of both tumor suppressor genes and oncogenes involved in CRC using data from existing databases and literature [60,76,98,100,101] against patient-specific mutations taken from cBioPortal [17]. The therapeutic screening was performed by upregulating the tumor suppressors and downregulating the oncogenes (Table S23), to evaluate potential drug combination targets using PanDrugs [77] database, a platform that prioritizes direct and indirect targeting of genomic mutations (see Supplementary Data 6).

### Combination of chemotherapy and targeted therapy to treat CRC patients

To induce the effect of chemotherapy we carried an extensive survey of the existing literature and constructed a microtubule network. The network consisted of 23 nodes and 28 edges. Next, this network was integrated with ISC-Apical network via up- and downstream signaling interaction. The resultant integrated network contained 39 nodes and 64 interactions. This integrated network was then utilized for chemotherapeutic screening. The combinatorial personalized therapy was used to treat the CRC patients, in a vertical therapy scheme through targeting specific nodes in our ISC-Apical network in light of patient mutations. DA pipeline was used to carry out the therapeutic evaluation (see Supplementary Data 6).

## Authors’ contributions

SUC designed and supervised the study. MNG carried out the literature review, construction of the model, and undertook the analyses. MNG and RNB designed the personalized treatment pipeline, SUC, MNG, RNB, ZN, and HK drafted the manuscript. OSS helped construct Boolean networks. RH critically reviewed the model development and performed validations, MT and AF assisted in the study design and manuscript development. All authors read and approved the final manuscript.

## Acknowledgments

This work was supported by the National ICT-R&D Fund (SRG-209), 15 NGIRI-2020-4771, HEC (21-30SRGP/R&D/HEC/ 2014, 20-2269/NRPU/R&D/ 16 HEC/12/ 4792 and 20-3629/NRPU/R&D/HEC/14/ 585), TWAS (RG 14-319 RG/ITC/AS_C) and LUMS (STG-BIO-1008, FIF-BIO-2052, FIF-BIO-0255, SRP-185-BIO, SRP-058-BIO and FIF-477-1819-BIO) grants.

## Supplementary material

**Supplementary Figure 1 Schematic representation of regulation in Intestinal Stem Cells**

Intestinal Stem Cell (ISC) in both apical and basal compartments employ six major signaling pathways including EGFR, WNT, JAK/STAT, BMP, NOTCH and Robo to maintain homeostasis and regeneration in the midgut. The inputs (green boxes) to these pathways are EGFs, Wg, Upds, Dpp, Delta and Slit, respectively. Each input is mapped to the output through an intermediate layer of nodes. The outputs (orange boxes) include Rolled, Cdc42, Hid, Dlg, Apc-Arm, TCF-LEF, STAT92E, Su(H), and Pros. The output layer is used to program cell fates which includes Extrusion, Apoptosis, Polarity, Division, Multilayering, Delta and Upd Production, EB Fate and EE Fates.

**Supplementary Figure 2 Schematic representation of regulation in Enteroblast**

Enteroblasts employ five major signaling pathways including EGFR, WNT, JAK/STAT, BMP, and NOTCH. The inputs (green boxes) to these pathways are EGFs, Wg, Upds, Dpp, and Delta, respectively. Each input is mapped on to the output through an intermediate layer of nodes. The outputs (orange boxes) include Cdc42, Hid, Dlg, Apc-Arm, STAT92E, and Su(H). The output layer is used to program cell fates which include Extrusion, Apoptosis, Polarity, Multilayering, Delta and Upd Production and EC Fate.

**Supplementary Figure 3 Schematic representation of regulation in Enterocyte**

Enterocytes employ four major signaling pathways including EGFR, WNT, BMP, and NOTCH. The inputs (green boxes) to these pathways are EGFs, Wg, Dpp, and Delta, respectively. Each input is mapped to the output through an intermediate layer of nodes. The outputs (orange boxes) include Cdc42, Hid, Dlg, Apc-Arm, and Su(H). The output layer is used to program cell fates which include Extrusion, Apoptosis, Polarity, Multilayering, Dpp and Upd Production.

**Supplementary Figure 4 Schematic representation of regulation in Enteroendocrine**

Enteroendocrine cells employ four major signaling pathways including EGFR, WNT, BMP, and NOTCH. The input (green boxes) to these pathways are EGFs, Wg, Dpp, and Delta, respectively. Each input is mapped to the output through an intermediate layer of nodes. The outputs (orange boxes) include Cdc42, Hid, Dlg, Apc-Arm, and Su(H). The output layer is used to program cell fates which include Extrusion, Apoptosis, Polarity, Multilayering, and Upd Production.

**Supplementary Figure 5 Schematic representation of regulation in Visceral Muscle cells**

Visceral Muscle cells employ five major signaling pathways including EGFR, WNT, JAK/STAT, BMP, and NOTCH. The inputs (green boxes) to these pathways are EGFs, Wg, Upds, Dpp, and Delta, respectively. Each input is mapped on to outputs through an intermediate layer of nodes. The outputs (orange boxes) include Cdc42, Hid, Dlg, Apc-Arm, STAT92E, and Su(H). The output layer is used to program cell fates which include Apoptosis, Wnt target genes, Delta, Upd and Dpp Production.

**Supplementary Figure 6 TISON implementation of biomolecular pathways involved in regulating intestinal stem cells (apical and basal)**

The network contains 32 nodes and 50 edges with 6 input, 9 output and 17 processing nodes.

**Supplementary Figure 7 TISON implementation of biomolecular pathways involved in regulating Enteroblast**

The network contains 29 nodes and 45 edges with 5 input, 6 output and 18 processing nodes.

**Supplementary Figure 8 TISON implementation of biomolecular pathways involved in regulating Enterocyte**

The network contains 23 nodes and 35 edges with 4 input, 6 output and 13 processing nodes.

**Supplementary Figure 9 TISON implementation of biomolecular pathways involved in regulating Enteroendocrine**

The network contains 23 nodes and 35 edges with 4 input, 6 output and 13 processing nodes.

**Supplementary Figure 10 TISON implementation of biomolecular pathways involved in regulating Visceral Muscle**

The network contains 26 nodes and 37 edges with 5 input, 5 output and 16 processing nodes.

**Supplementary Figure 11 Standard Error of Means (SEM) for ISC, EB, EC and VM**

The SEM with highest for Apoptosis in ISC (0.00128), Upd production for EB (0.00287), Dpp Production for EC (0.0026) and WNT target gene fate for VM 15 (0.0039).

**Supplementary Figure 12 Circos plot of biomolecular regulatory networks in ISC (purple), EB (blue), EC (orange), EE (red) and VM (green) cells**

The plots shows the interaction relationship between 5 cell types, 12 cell fates and their respective pathways.

**Supplementary Figure 13 Intestinal Stem Cell – Apical, in Normal Condition**

**Supplementary Figure 14 Intestinal Stem Cell – Apical, in Stress Condition**

**Supplementary Figure 15 Intestinal Stem Cell – Apical, in Cancer Condition**

**Supplementary Figure 16 Intestinal Stem Cell – Basal, in Normal Condition**

**Supplementary Figure 17 Intestinal Stem Cell – Basal, in Stress Condition**

**Supplementary Figure 18 Intestinal Stem Cell – Basal, in Cancer Condition**

**Supplementary Figure 19 Enteroblast, in Normal Condition**

**Supplementary Figure 20 Enteroblast, in Stress Condition**

**Supplementary Figure 21 Enteroblast, in Cancer Condition**

**Supplementary Figure 22 Enterocyte, in Normal Condition**

**Supplementary Figure 23 Enterocyte, in Stress Condition**

**Supplementary Figure 24 Enterocyte, in Cancer Condition**

**Supplementary Figure 25 Schematic of homeostasis, differentiation and tumorigenesis in normal and diseased midgut**

**A**. In normal midgut, basal ISCs maintain stemness or differentiate into EE while apical ISCs get converted into EB. EBs can then differentiate into ECs under certain conditions; however, they mostly remain dormant in homeostatic conditions. **B**. In diseased midgut, depending on the mutation type, the gut can either form adenocarcinoma or carcinoma. APC mutation can lead to development of an adenocarcinoma in the gut with a further Ras mutation can result in carcinoma.

**Supplementary Figure 26 Schematic Representation of Regulation in Mitochondria**

The inputs (green boxes) to these pathways are GF, Upds, Slit and Wg, respectively. Each input is mapped to the output through an intermediate layer of nodes. The outputs (orange boxes) include Stathmin, CLASP, CDK, and Apc-Arm. The output layer is used to program cell fates which includes Destabilize and Stabilize microtubule, corresponding to Proliferation and Apoptosis cell fates, respectively.

**Supplementary Figure 27 Schematic Representation of Regulation an Integrated Network of Intestinal Stem Cells and Microtubule**

The inputs (green boxes) to these pathways are GF, EGFs, Wg, Upds, Dpp, Delta, and Slit, respectively. Each input is mapped to the output through an intermediate layer of nodes. The outputs (orange boxes) include Rolled, Cdc42, Hid, Dlg, Apc-Arm, TCF-LEF, STAT92E, Su(H), Stathmin, CLASP, CDK and Pros. The output layer is used to program cell fates which includes Extrusion, Apoptosis, Polarity, Division, Multilayering, Delta and Upd Production, EB Fate and EE Fates.

**Supplementary Table 1 Detailed node interaction rules and experimental evidences supporting different interactions and logical functions for Intestinal Stem Cells (ISC) model**

**Supplementary Table 2 Detailed node interaction rules and experimental evidences supporting different interactions and logical functions for Enteroblast (EB) model**

**Supplementary Table 3 Detailed node interaction rules and experimental evidences supporting different interactions and logical functions for Enterocyte (EC) model**

**Supplementary Table 4 Detailed node interaction rules and experimental evidences supporting different interactions and logical functions for Enteroendocrine (EE) model**

**Supplementary Table 5 Detailed node interaction rules and experimental evidences supporting different interactions and logical functions for Visceral Muscle (VM) cells model**

**Supplementary Table 6 Robustness analysis cell fates and corresponding SEMs for ISC, EB, EC and VM**

**Supplementary Table 7 Fixed node input states in normal, stress and cancer for ISC-Apical, ISC-Basal, EB and EC network models and their literature validation**

**Supplementary Table 8 Cell fate propensities of Intestinal Stem Cells (ISC) in Apical and Basal compartments; Enteroblast (EB) and Enterocytes (EC) in Normal, Stress and Cancer conditions along with literature validations**

**Supplementary Table 9 A comparison of model and experimental output node propensities**

**Supplementary Table 10 Tabulation of network nodes, gene IDs, annotation symbols, gene symbols, and FlyBase genes**

**Supplementary Table 11 Martorell *et al’*s predictions: experiment versus model**

**Supplementary Table 12 Differential gene expression comparison between prediction and the model**

**Supplementary Table 13 Results of class I and class II drugs from Markstein *et al*.’s therapeutics screens**

**Supplementary Table 14 Cell fate propensities for proliferation and apoptosis in class I and class II drugs**

**Supplementary Table 15 Details of the Bangi *et al*.’s case study: mutations, therapy, and induction of therapy in the *in silico* DPM**

**Supplementary Table 16 Oncogenic cell fate propensities of potential target nodes**

**Supplementary Table 17 Personalized therapeutic combinations for individual patients**

**Supplementary Table 18 Detailed node interaction rules and experimental evidences supporting different interactions and logical functions Microtubule model (MT)**

**Supplementary Table 19 Detailed node interaction rules and experimental evidences supporting different interactions and logical functions for Integrated (ISC+MT) model**

**Supplementary Table 20 Results from chemotherapy + targeted therapy of colorectal cancer patients**

**Supplementary Table 21 Results from targeted therapy of colorectal cancer patients**

**Supplementary Table 22 Mapping of cell fate classification logic**

**Supplementary Table 23 Details of tumor suppressors and oncogenes in ISC-Apical network**

## Supplementary URL link

The supplementary data of manuscript titled “*In silico Drosophila Patient Model Reveals Optimal Combinatorial Therapies for Colorectal Cancer*” including input files, output files and analysis results are available at GitHub on this URL: https://github.com/BIRL/DrosophilaPatientModel

## References

1. Hanahan D, Weinberg RA. Hallmarks of Cancer: The Next Generation. Cell. 2011 Mar;144(5):646–74.

2. Baylin SB, Jones PA. A decade of exploring the cancer epigenome — biological and translational implications. Nat Rev Cancer [Internet]. 2011;11(10):726–34. Available from: https://doi.org/10.1038/nrc3130

3. Bild AH, Yao G, Chang JT, Wang Q, Potti A, Chasse D, et al. Oncogenic pathway signatures in human cancers as a guide to targeted therapies. Nature. 2006;439(7074):353–7.

4. Harris CC, Hollstein M. Clinical implications of the p53 tumor-suppressor gene. N Engl J Med. 1993;329(18):1318–27.

5. Mirzoyan Z, Sollazzo M, Allocca M, Valenza AM, Grifoni D, Bellosta P. Drosophila melanogaster: A model organism to study cancer. Front Genet. 2019;10(March):1–16.

6. Bujanda L, Cosme A, Gil I, Arenas-Mirave JI. Malignant colorectal polyps. Vol. 16, World journal of gastroenterology. 2010. p. 3103–11.

7. Hanahan D, Weinberg RA, Francisco S. The Hallmarks of Cancer Review University of California at San Francisco. 2000;100:57–70.

8. Meacham CE, Morrison SJ. Tumour heterogeneity and cancer cell plasticity. Vol. 501, Nature. Howard Hughes Medical Institute; 2013. p. 328–37.

9. Dagogo-Jack I, Shaw AT. Tumour heterogeneity and resistance to cancer therapies. Vol. 15, Nature Reviews Clinical Oncology. Nature Publishing Group; 2018. p. 81–94.

10. Szakács G, Paterson JK, Ludwig JA, Booth-Genthe C, Gottesman MM. Targeting multidrug resistance in cancer. Nat Rev Drug Discov. 2006;5(3):219–34.

11. Ocana A, Pandiella A, Siu LL, Tannock IF. Preclinical development of molecular-targeted agents for cancer. Nat Rev Clin Oncol [Internet]. 2011;8(4):200–9. Available from: https://doi.org/10.1038/nrclinonc.2010.194

12. DiMasi JA, Reichert JM, Feldman L, Malins A. Clinical approval success rates for investigational cancer drugs. Clin Pharmacol Ther. 2013;94(3):329–35.

13. Zamboni WC, Torchilin V, Patri AK, Hrkach J, Stern S, Lee R, et al. Best practices in cancer nanotechnology: perspective from NCI nanotechnology alliance. lin cancer Res. 2012;18(12):3229–41.

14. Gottesman MM, Fojo T, Bates SE. Multidrug resistance in cancer: role of ATP–dependent transporters. Nat Rev Cancer. 2002;2(1):48–58.

15. Kasai Y, Cagan R. Drosophila as a tool for personalized medicine: a primer. Per Med. 2010;7(6):621–32.

16. Grandori C, Kemp CJ. Personalized cancer models for target discovery and precision medicine. Trends in cancer. 2018;4(9):634–42.

17. Gao J, Aksoy BA, Dogrusoz U, Dresdner G, Gross B, Sumer SO, et al. Integrative analysis of complex cancer genomics and clinical profiles using the cBioPortal. Sci Signal. 2013 Apr;6(269):11.

18. Lee HJ, Palm J, Grimes SM, Ji HP. The Cancer Genome Atlas Clinical Explorer: A web and mobile interface for identifying clinical-genomic driver associations. Genome Med [Internet]. 2015;7(1):1–14. Available from: 19 http://dx.doi.org/10.1186/s13073-015-0226-3

19. Zhang J, Baran J, Cros A, Guberman JM, Haider S, Hsu J, et al. International Cancer Genome Consortium Data Portal—a one-stop shop for cancer genomics data. Database. 2011;2011.

20. Zhang L, Jiang B, Wu Y, Strouthos C, Sun PZ, Su J, et al. Developing a multiscale, multi-resolution agent-based brain tumor model by graphics processing units. Theor Biol Med Model [Internet]. 2011;8(1):46. Available from: https://doi.org/10.1186/1742-4682-8-46

21. Li J, Akbani R, Zhao W, Lu Y, Weinstein JN, Mills GB, et al. Explore, visualize, and analyze functional cancer proteomic data using the cancer proteome atlas. Cancer Res. 2017;77(21):e51–4.

22. Thul PJ, Lindskog C. The human protein atlas: A spatial map of the human proteome. Protein Sci. 2018;27(1):233–44.

23. Morgan MM, Johnson BP, Livingston MK, Schuler LA, Alarid ET, Sung KE, et al. Personalized in vitro cancer models to predict therapeutic response: Challenges and a framework for improvement. Pharmacol Ther. 2016;165:79–92.

24. Lieschke GJ, Currie PD. Animal models of human disease: zebrafish swim into view. Nat Rev Genet. 2007;8(5):353.

25. Olive KP, Tuveson DA. The use of targeted mouse models for preclinical testing of novel cancer therapeutics. Clin Cancer Res. 2006;12(18):5277–87.

26. Pandey UB, Nichols CD. Human Disease Models in Drosophila melanogaster and the Role of the Fly in Therapeutic Drug Discovery. 2011;63(2):411–36.

27. Adams MD, Celniker SE, Holt RA, Evans CA, Gocayne JD, Amanatides PG, et al. The genome sequence of Drosophila melanogaster. Science (80-). 2000;287(5461):2185–95.

28. Reiter LT, Potocki L, Chien S, Gribskov M, Bier E. A systematic analysis of human disease-associated gene sequences in Drosophila melanogaster. Genome Res. 2001 Jun;11(6):1114–25.

29. Rubin GM, Yandell MD, Wortman JR, Gabor Miklos GL, Nelson CR, Hariharan IK, et al. Comparative genomics of the eukaryotes. Science. 2000 Mar;287(5461):2204–15.

30. Drysdale RA, Consortium F, Crosby MA, Consortium F. FlyBase: genes and gene models. Nucleic Acids Res. 2005;33(suppl_1):D390–5.

31. Buchon N, Edgar B. Flygut-seq: Cell and region specific gene expression of the fly midgut [Internet]. Available from: http://flygutseq.buchonlab.com/references

32. Chintapalli VR, Wang J, Dow JAT. Using FlyAtlas to identify better Drosophila melanogaster models of human disease. Nat Genet. 2007;39(6):715–20.

33. Vidal M, Wells S, Ryan A, Cagan R. ZD6474 suppresses oncogenic RET isoforms in a Drosophila model for type 2 multiple endocrine neoplasia syndromes and papillary thyroid carcinoma. Cancer Res. 2005;65(9):3538–41.

34. Sonoshita M, Cagan RL. Modeling Human Cancers in Drosophila [Internet]. 1st ed. Vol. 121, Fly Models of Human Diseases. Elsevier Inc.; 2016. 287–309 p. Available from: http://dx.doi.org/10.1016/bs.ctdb.2016.07.008

35. Bangi E, Garza D, Hild M. In vivo analysis of compound activity and mechanism of action using epistasis in Drosophila. J Chem Biol. 2011;4(2):55–68.

36. Levine BD, Cagan RL. Drosophila lung cancer models identify trametinib plus statin as candidate therapeutic. Cell Rep. 2016;14(6):1477–87.

37. Bossen J, Uliczka K, Steen L, Pfefferkorn R, Mai MM-Q, Burkhardt L, et al. An EGFR-induced Drosophila lung tumor model identifies alternative combination treatments. Mol Cancer Ther. 2019;18(9):1659–68.

38. Bhandari P, Shashidhara LS. Studies on human colon cancer gene APC by targeted expression in Drosophila. Oncogene. 2001;20(47):6871–80.

39. Palmer AC, Sorger PK. Combination Cancer Therapy Can Confer Benefit via Patient-to-Patient Variability without Drug Additivity or Synergy. Cell [Internet]. 2017 Dec 14;171(7):1678–1691.e13. Available from: https://pubmed.ncbi.nlm.nih.gov/29245013

40. Chen S-H, Lahav G. Two is better than one; toward a rational design of combinatorial therapy. Curr Opin Struct Biol [Internet]. 2016/08/10. 2016 Dec;41:145–50. Available from: https://pubmed.ncbi.nlm.nih.gov/27521655

41. Yardley DA. Drug Resistance and the Role of Combination Chemotherapy in Improving Patient Outcomes. Fentiman IS, editor. Int J Breast Cancer [Internet]. 2013;2013:137414. Available from: https://doi.org/10.1155/2013/137414

42. Aird RE, Cummings J, Ritchie AA, Muir M, Morris RE, Chen H, et al. In vitro and in vivo activity and cross resistance profiles of novel ruthenium (II) organometallic arene complexes in human ovarian cancer. Br J Cancer. 2002 May;86(10):1652–7.

43. Murayama T, Gotoh N. Patient-derived xenograft models of breast cancer and their application. Cells. 2019;8(6):621.

44. Bangi E, Ang C, Smibert P, Uzilov A V, Teague AG, Antipin Y, et al. A personalized platform identifies trametinib plus zoledronate for a patient with KRAS-mutant metastatic colorectal cancer. Sci Adv. 2019;5(5):eaav6528.

45. Passini E, Britton OJ, Lu HR, Rohrbacher J, Hermans AN, Gallacher DJ, et al. Human In Silico Drug Trials Demonstrate Higher Accuracy than Animal Models in Predicting Clinical Pro-Arrhythmic Cardiotoxicity. Front Physiol [Internet]. 2017;8:668. Available from: https://www.frontiersin.org/article/10.3389/fphys.2017.00668

46. Richardson HE, Willoughby L, Humbert PO. Screening for Anti-cancer Drugs in Drosophila. eLS. 2001;1–14.

47. Micchelli CA, Perrimon N. Evidence that stem cells reside in the adult Drosophila midgut epithelium. Nature. 2006;439(7075):475.

48. Ohlstein B, Spradling A. The adult Drosophila posterior midgut is maintained by pluripotent stem cells. Nature. 2006;439(7075):470.

49. Takashima S, Adams KL, Ortiz PA, Ying CT, Moridzadeh R, Younossi-Hartenstein A, et al. Development of the Drosophila entero-endocrine lineage and its specification by the Notch signaling pathway. Dev Biol. 2011;353(2):161–72.

50. Ohlstein B, Spradling A. Multipotent Drosophila intestinal stem cells specify daughter cell fates by differential notch signaling. Science (80-). 2007;315(5814):988–92.

51. Lee M, Vasioukhin V. Cell polarity and cancer – cell and tissue polarity as a non-canonical tumor suppressor. 2008;

52. Pasco MY, Loudhaief R, Gallet A. The cellular homeostasis of the gut : what the Drosophila model points out. 2014;(October).

53. Loza-coll MA, Southall TD, Sandall SL, Brand AH, Jones DL. Regulation of Drosophila intestinal stem cell maintenance and differentiation by the transcription factor Escargot. 2014;33(24):2983–96.

54. Zeng X, Hou SX. Enteroendocrine cells are generated from stem cells through a distinct progenitor in the adult Drosophila posterior midgut. Development [Internet]. 2015;142(4):644–53. Available from: http://dev.biologists.org/cgi/doi/10.1242/dev.113357

55. Macara IG, McCaffrey L. Cell polarity in morphogenesis and metastasis. Philos Trans R Soc London Ser B, Biol Sci. 2013 Nov;368(1629):20130012.

56. Bidirectional Notch signaling regulates Drosophila intestinal stem cell multipotency. 2015;350(6263).

57. Snoeck V, Goddeeris B, Cox E. The role of enterocytes in the intestinal barrier function and antigen uptake. 2005;7:997–1004.

58. Wolfstetter G, Shirinian M, Stute C, Grabbe C, Hummel T, Baumgartner S, et al. Fusion of circular and longitudinal muscles in Drosophila is independent of the endoderm but further visceral muscle differentiation requires a close contact between mesoderm and endoderm. Mech Dev [Internet]. 2009;126(8):721–36. Available from: http://www.sciencedirect.com/science/article/pii/S092547730900063X

59. FlyGut-seq.

60. Martorell Ò, Merlos-Suárez A, Campbell K, Barriga FM, Christov CP, Miguel-Aliaga I, et al. Conserved mechanisms of tumorigenesis in the Drosophila adult midgut. PLoS One. 2014;9(2).

61. Markstein M, Dettorre S, Cho J, Neumuller RA, Craig-Muller S, Perrimon N. Systematic screen of chemotherapeutics in Drosophila stem cell tumors. Proc Natl Acad Sci [Internet]. 2014;111(12):4530–5. Available from: http://www.pnas.org/cgi/doi/10.1073/pnas.1401160111

62. cBioPortal for Cancer Genomics.

63. Stone L. The feasibility and stability of large complex biological networks: a random matrix approach. Sci Rep [Internet]. 2018 May 29;8(1):8246. Available from: https://pubmed.ncbi.nlm.nih.gov/29844420

64. Kwon Y-K, Cho K-H. Quantitative analysis of robustness and fragility in biological networks based on feedback dynamics. Bioinformatics [Internet]. 2008;24(7):987–94. Available from: https://doi.org/10.1093/bioinformatics/btn060

65. Glass L, Kauffman SA. The logical analysis of continuous, non-linear biochemical control networks. J Theor Biol. 1973 Apr;39(1):103–29.

66. Amcheslavsky A, Jiang J, Ip YT. Tissue damage-induced intestinal stem cell division in Drosophila. Cell Stem Cell. 2009 Jan;4(1):49–61.

67. Apidianakis Y, Pitsouli C, Perrimon N, Rahme L. Synergy between bacterial infection and genetic predisposition in intestinal dysplasia. Proc Natl Acad Sci U S A. 2009 Dec;106(49):20883–8.

68. Schell JC, Wisidagama DR, Bensard C, Zhao H, Wei P, Tanner J, et al. Control of intestinal stem cell function and proliferation by mitochondrial pyruvate metabolism. Nat Cell Biol. 2017 Sep;19(9):1027–36.

69. Di Biase S, Longo VD. Fasting-induced differential stress sensitization in cancer treatment. Mol Cell Oncol. 2016 May;3(3):e1117701.

70. Adlesic M, Frei C, Frew IJ. Cdk4 functions in multiple cell types to control Drosophila intestinal stem cell proliferation and differentiation. Biol Open. 2016 Feb;5(3):237–51.

71. O’Brien LE, Soliman SS, Li X, Bilder D. Altered modes of stem cell division drive adaptive intestinal growth. Cell. 2011 Oct;147(3):603–14.

72. Hogan C, Dupré-Crochet S, Norman M, Kajita M, Zimmermann C, Pelling AE, et al. Characterization of the interface between normal and transformed epithelial cells. Nat Cell Biol [Internet]. 2009 Mar 15;11:460. Available from: https://doi.org/10.1038/ncb1853

73. Sancho E, Batlle E, Clevers H. SIGNALING PATHWAYS IN INTESTINAL DEVELOPMENT AND CANCER. Annu Rev Cell Dev Biol [Internet]. 2004;20(1):695–723. Available from: https://doi.org/10.1146/annurev.cellbio.20.010403.092805

74. Network TCGA, Muzny DM, Bainbridge MN, Chang K, Dinh HH, Drummond JA, et al. Comprehensive molecular characterization of human colon and rectal cancer. Nature [Internet]. 2012 Jul 18;487:330. Available from: https://doi.org/10.1038/nature11252

75. Oberley LW, Oberley TD, Buettner GR. Cell differentation, aging and cancer: The possible roles of superoxide and superoxide dismutases. Med Hypotheses [Internet]. 1980;6(3):249–68. Available from: http://www.sciencedirect.com/science/article/pii/0306987780901231

76. Aleman M. Modelling colorectal cancer in Drosophila.

77. Piñeiro-Yáñez E, Reboiro-Jato M, Gómez-López G, Perales-Patón J, Troulé K, Rodríguez JM, et al. PanDrugs: a novel method to prioritize anticancer drug treatments according to individual genomic data. Genome Med. 2018;10(1):1–11.

78. Rowinsky EK, Donehower RC. Paclitaxel (taxol). N Engl J Med. 1995;332(15):1004–14.

79. Han ES, Wen W, Dellinger TH, Wu J, Lu SA, Jove R, et al. Ruxolitinib synergistically enhances the anti-tumor activity of paclitaxel in human ovarian cancer. Oncotarget. 2018;9(36):24304.

80. Fruehauf JP, El-Masry M, Osann K, Parmakhtiar B, Yamamoto M, Jakowatz JG. Phase II study of pazopanib in combination with paclitaxel in patients with metastatic melanoma. Cancer Chemother Pharmacol. 2018 Aug;82(2):353–60.

81. Pazopanib and Paclitaxel for Non-Small Cell Lung Cancer [Internet]. 2010. Available from: https://clinicaltrials.gov/ct2/show/NCT01179269

82. Pink D, Bauer S, Brodowicz T, Reichardt P, Kasper B, Richter S, et al. Treatment of angiosarcoma with pazopanib and paclitaxel: Results of the phase II trial of the German Interdisciplinary Sarcoma Group (GISG-06 EVA) study. J Clin Oncol [Internet]. 2018 May 20;36(15_suppl):11570. Available from: https://doi.org/10.1200/JCO.2018.36.15_suppl.11570

83. Kanehisa M. KEGG: Kyoto Encyclopedia of Genes and Genomes. Nucleic Acids Res. 2000 Jan;28(1):27–30.

84. Yu J, Pacifico S, Liu G, Finley RL. DroID: the Drosophila Interactions Database, a comprehensive resource for annotated gene and protein interactions. BMC Genomics. 2008;9(1):1–9.

85. Gelbart WM, Rindone WP, Chillemi J, Russo S, Crosby M, Mathews B, et al. FlyBase: The Drosophila database. Vol. 24, Nucleic Acids Research. 1996. p. 53–6.

86. Giot L, Bader JS, Brouwer C, Chaudhuri A, Kuang B, Li Y, et al. A Protein Interaction Map of Drosophila melanogaster. Science (80-). 2003;302(5651):1727–36.

87. Formstecher E, Aresta S, Collura V, Hamburger A, Meil A, Trehin A, et al. Protein interaction mapping: A Drosophila case study. Genome Res. 2005;15(3):376–84.

88. Vinson KE, George DC, Fender AW, Bertrand FE, Sigounas G. The Notch pathway in colorectal cancer. 2016;

89. Buchon N, Broderick NA, Kuraishi T, Lemaitre B. Drosophila EGFR pathway coordinates stem cell proliferation and gut remodeling following infection. BMC Biol. 2010;8(1):152.

90. Xu N, Wang SQ, Tan D, Gao Y, Lin G, Xi R. EGFR, Wingless and JAK/STAT signaling cooperatively maintain Drosophila intestinal stem cells. Dev Biol. 2011;354(1):31–43.

91. Slattery ML, Lundgreen A, Kadlubar SA, Bondurant KL, Wolff RK. JAK/STAT/SOCS-signaling pathway and colon and rectal cancer. Mol Carcinog. 2013 Feb;52(2):155–66.

92. Huang T, Kang W, Cheng ASL, Yu J, To KF. The emerging role of Slit-Robo pathway in gastric and other gastro intestinal cancers. BMC Cancer. 2015;15:950.

93. Glass L, Kauffman SA. The Logical Analysis of Continuous, Non-linear BiochemicaldaControl Networks. 1973;103–29.

94. Darabos C, Cunto F Di, Tomassini M, Moore JH, Provero P. Additive Functions in Boolean Models of Gene Regulatory Network Modules. 2011;6(11).

95. Shaun J. Grannis MD, J. Marc Overhage MD PhD CJMM. Analysis of identifier performance using a deterministic linkage algorithm. Proc AMIA Symp. 2002;305–9.

96. Shah OS, Chaudhary MFA, Awan HA, Fatima F, Arshad Z, Amina B, et al. ATLANTIS – Attractor Landscape Analysis Toolbox for Cell Fate Discovery and Reprogramming. Sci Rep [Internet]. 2018;8(1):1–11. Available from: http://dx.doi.org/10.1038/s41598-018-22031-3

97. Buchon N, Osman D, David FPA, Yu Fang H, Boquete J-P, Deplancke B, et al. Morphological and Molecular Characterization of Adult Midgut Compartmentalization in Drosophila. Cell Rep [Internet]. 2013;3(5):1725–38. Available from: http://www.sciencedirect.com/science/article/pii/S221112471300168X

98. Smirnov P, Kofia V, Maru A, Freeman M, Ho C, El-Hachem N, et al. PharmacoDB: an integrative database for mining in vitro anticancer drug screening studies. Nucleic Acids Res. 2018;46(D1):D994–1002.

99. Griffith M, Griffith OL, Coffman AC, Weible J V, McMichael JF, Spies NC, et al. DGIdb: mining the druggable genome. Nat Methods. 2013;10(12):1209–10.

100. Yang W, Soares J, Greninger P, Edelman EJ, Lightfoot H, Forbes S, et al. Genomics of Drug Sensitivity in Cancer (GDSC): A resource for therapeutic biomarker discovery in cancer cells. Nucleic Acids Res. 2013;41(D1).

101. Wishart DS, Feunang YD, Guo AC, Lo EJ, Marcu A, Grant JR, et al. DrugBank 5.0: a major update to the DrugBank database for 2018. Nucleic Acids Res. 2018;46(D1):D1074–82.

